# 3D-printable tools for developmental biology: Improving embryo injection and screening techniques through 3D-printing technology

**DOI:** 10.1101/376657

**Authors:** Marta Truchado-Garcia, Richard M. Harland, Michael J. Abrams

**Affiliations:** Departamento de Biología, Universidad Autónoma de Madrid, Calle Darwin, 2. Edificio de Biología. 28049 Madrid, Spain; Department of Molecular and Cell Biology, University of California, Berkeley, Life Sciences Addition #3200, Berkeley, CA 94720-3200, USA; Division of Biology and Biological Engineering, California Institute of Technology, Pasadena, CA 91125, USA

**Author notes:** **Corresponding authors:** Marta Truchado-Garcia, (new location) Department of Molecular and Cell Biology, University of California, Berkeley, Life Sciences Addition #3200, Berkeley, CA 94720-3200, USA. Michael J. Abrams, Division of Biology and Biological Engineering, California Institute of Technology, Pasadena, CA 91125, USA. Co-authors. **Competing interests:** The authors have declared that no competing interests exist.

**Keywords:** 3D-printing tools, embryos, injection, microscopy, screening

## Abstract

Developmental biology requires rapid embryo injections and screening. We applied new affordable high-resolution 3D-printing to create <u>five</u> easily modifiable stamp-mold tools that greatly increase injection and screening speed, while simultaneously reducing the harmful aspects of these processes. We designed two stamps that use different approaches to improve the injection efficiency for two different types of embryo, first for embryos from the snail *Crepidula fornicata*, and second, for those from the spider *Parasteatoda tepidariorum*. Both drastically improved injection speeds and embryo survival rates, even in novice hands. The other three tools were designed for rapid side-by-side organism orientating and comparison. The first screening tool allows for optimal imaging in *Xenopus laevis* tadpoles, while the second and third facilitate rapid high-throughput screening of *Xenopus tropicalis* tadpoles and *Danio rerio* juveniles, respectively. These designs can act as templates for many injection or screening applications.

## INTRODUCTION

Sometimes no protocol exists to answer the pertinent question, and we must creatively combine techniques and invent new ones. Sometimes the technique is quite difficult; in order to understand how embryos reorganize their cells during gastrulation, Keller and Danilchik [1, 2] developed a clever technique to use on *Xenopus* embryos called “the sandwich,” consisting of removing strips of dorsal mesendoderm and ectoderm from embryos in early gastrulae with a eye lash and squeezing it into clay at the bottom of a petri dish in specific orientations. But even simple and common tasks can be very tedious and take considerable engineering work, and countless hours of trial and error. With modern molecular biology, the recently almost unimaginable act of sequencing or cloning DNA is now easier to perform than some embryological experiments that have been common practice for decades.

Embryo microinjections and screening are essential techniques in developmental biology that are routinely practiced. In order to successfully accomplish a comprehensive project, lineage tracing and transgenesis experiments are often performed. These studies generally require the injection of reagents such as dextran, diI, morpholinos, RNAi, mRNA *in vitro* synthesized or constructs inside vectors [3, 4, 5, 6]. Subsequently, checking the embryos, or the frequently highly mobile juveniles, is an essential next step for correctly describing a phenotype or a whole *in vivo* process [7, 8]. Generally, each individual laboratory has its own special injection and screening system. For example, some laboratories make customized injection apparatuses out of glass covers and slides [9], while others scratch plastic dishes to generate sticky edges [8]. For screening, larvae are often placed in relatively toxic compounds [10] or require tedious procedures with methyl-cellulose, low-melting point agarose [11] or heptane-glue mixtures for immobilization [12]. These routine practices require many months of training and practice to master and can be harmful to the organisms. In zebrafish, injection of the zygotes and specific cells is a routine task and this model, as well as *Xenopus*, have the advantage of producing big zygotes (0.7 mm in diameter; [13]), and yet their developing progeny begin to wriggle making screening and long-term developmental imaging quite difficult.

Developmental biologists customize all kinds of tools to succeed at these protocols, including specialized forceps, stretched glass needles, miniature scalpels, and non-toxic sealants like Valap [14]. However, when making tools, size and resolution limitations have always established a barrier. We take advantage of new high-resolution 3D-printing technology that has recently become affordable enough for use in academia [15, 16]. Using free design software, and a relatively inexpensive high-precision stereolithographic 3D-printer (Form2), we developed two injection tools (a cup-tool for spider embryos and a trench-tool for snail embryos) that immobilize embryos without damaging them, even allowing novices to successfully inject difficult embryos. We also designed and tested screening tools that allow Xenopus and Zebrafish to be rapidly oriented for imaging. These stamp-mold designs can be customized to keep embryos and juveniles of practically any size or shape in their proper orientations for different applications.

## MATERIALS AND METHODS

### 3D-modeling and printing

Autodesk 123D (http://www.123dapp.com/design) was used to design the stamps as it was available for free. It has a helpful tutorial for beginners and has an intuitive interface.

All 3D-printing was performed using the FormLabs Form2 (http://formlabs.com/products/3d-printers/form-2/). Autodesk 123D designs were imported and prepared for printing using the FormLabs printing software named PreForm (http://formlabs.com/support/software/install-preform-software/). 3D-printing resolution is application dependent; however, we used the 0.025mm settings.

Using the FormLabs software to properly orient the stamp design for printing is important for optimal printing performance. The stamp should be oriented so that the front surface faces away from the 3D-printer’s printing surface, so supports that attach to the printing surface are not placed on the stamp face. Also, the stamp face should be at a 45-degree angle to the printing surface, with one of the stamp’s corners, and not the flat edge, closest to the printing surface. This allows for optimal printing resolution.

### Injection Tools: Designing the Injection Stamps and Molds

We developed one trough-style injection stamp for the snail *Crepidula fornicata*, which has zigotes that are 200 μM in diameter [8], and one cup-style for the spider *Parasteatoda tepidariorum*, which has zigotes that are 400 μm in diameter [17]. To prepare the injection mold, we use agarose 2% (agarose D-1 Low EEO, Pronadisa, Laboratorios Conda) diluted in artificial filtered seawater (AFSW) [8]. This agarose solution is poured into a p60 petri dish, and the stamp placed on the newly poured liquid agarose. It is important to consider the distance between the rim and the agarose surface, so that the needle does not break on the rim. Once it has slowly cooled down and solidified, the stamp is removed and the mold covered with FSW for injecting *Crepidula fornicata* embryos, and its embryos gently arranged inside the troughs and injected. For spiders, the mold generates embryo holder spaces, where they are loaded into the dish and adjusted into the spaces. The mold contains 126 wells (14 x 9).

### Designing the Screening Stamps and Molds

We designed one stamp to screen the tadpoles of *Xenopus laevis* in stage 43 and another for Zebrafish at 10 days after fertilization (after hatching; swimming juveniles).

Swimming tadpoles need to be anesthetized with MS-222 [7, 10]. The two by two-sided mold for Xenopus is generated by firmly pressing against the flattened modeling clay (wax based clay) in a p35 petri dish. Tadpoles grew up to stage 43 and were used for imaging. We anesthetized the animals directly on the dish, where they were oriented and imaged. Then we recovered the tadpoles on a rocker in a new dish [7]. For a more high-throughput screening process, a multi-mold *Xenopus tropicalis* tadpole stamp was designed, to simultaneously screen 20 (4 x 5) tadpoles, in a drastically more efficient process. The mold is made with agarose in the same way as the injection systems are prepared.

Similarly, the zebrafish screening stamp allows for 24 dorsal/ventral and 24 right/left zebrafish, side-by-side, to be analyzed at once, ideal for confocal microscopy. Inverted confocal LSM800 (Zeiss) was used in our description. A glass bottom dish was made by removing the bottom of a p35 dish and fixing a 24 x 24 coverslip to cover the space. We add 1 ml of agarose (agarose D-1 Low EEO, Pronadisa, Laboratorios Conda), to make it as transparent and thin as possible. The depth of the agarose should be shallow enough to allow the stamp shapes to be seen against the glass, and the fish should nearly touch the glass. E3 medium with tricaine is poured on the solidified agarose in order to anesthetize the animals. After the dish is put on the confocal platform, the fish were orientated with a lash-tool, and then imaged. The mutant line used expresses Patched-GFP and was kindly provided by Dr. Citlali Gradilla at Centro de Biología Molecular Severo-Ochoa (Madrid, Spain).

## RESULTS

### Injection stamp for *Crepidula* increases speed and survivability of embryos and larvae

The snail injection stamp (**Fig. 1C**) generates a mold (**Fig. 1B**) that has two injection troughs (**Fig. 1A-D**). The embryos are loaded into the mold-dish and adjusted into the troughs. The backstop prevents the embryo from being pushed away by the needle, while the troughs keep the embryos in a straight line for rapid injections (**Fig. 1A**). The combination of the troughs and the backstops allows this snail injection device to keep the embryos from moving without damaging them, the first important requirement for improved injections. If an embryo sticks to the injection needle, which is common, by simply pulling the needle back through the front-stop of the trough (**Fig. 1A and 1D**), the embryo can be gently squeezed off the needle tip, where it will remain in the trough, meeting a second important requirement [similar requirements for *Ciona intestinalis;* data not shown].

**Figure 1.**
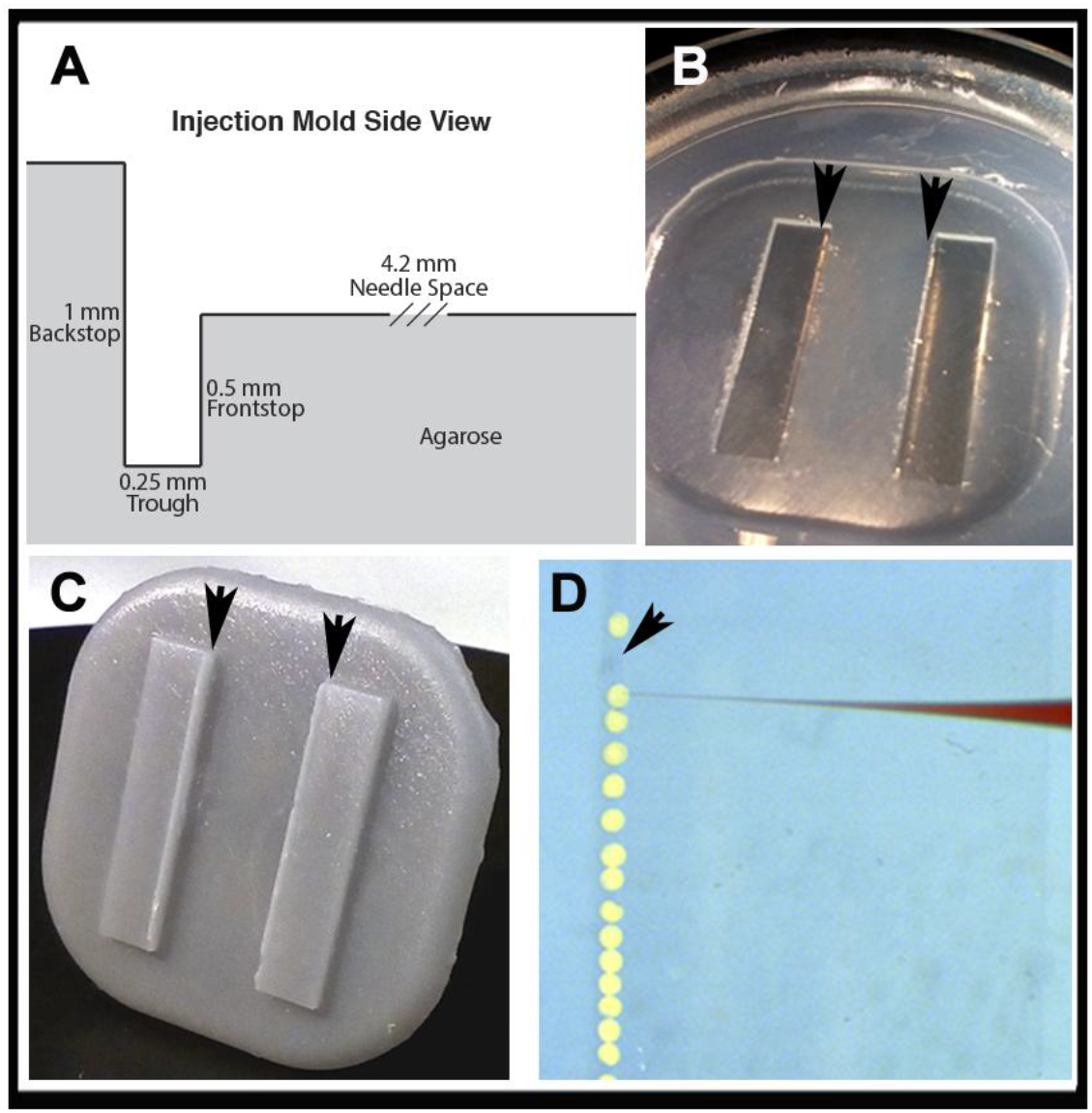
Injection stamp for *Crepidula fonicata* zygotes. **A.** Schematic of injection stamp-mold with labeled parameters. **B.** Agarose gel mold of injection stamp in a p60 petri dish, with arrows indicating the trench location. **C.** 3D-printed injection stamp showing the extruded needle spaces and troughs (arrows). **D.** *Crepidula fornicata* zygotes in the trough (arrow), with a loaded needle prepared for injections.

In standard practice *Crepidula* embryos are placed in a razor-made groove in a petri-dish (described in Discussion). In expert hands it is possible to inject about ~250 *Crepidula* embryos/hour, with over 95% survival, using that setup. However, it is common for novices injecting in this manner to get closer to 45 embryos successfully injected in the same amount of time, with a 58% survival rate.

Startlingly, our injection system has allowed complete novices, just over the course of one day of injections, to drastically improve their efficiency. Novices have increased the number of injected embryos to 80 in the same set of experiments, and even more, increased the survival rate up ~70%. This addresses many of the previously described problems, allowing novices to quickly gather more data in an efficient and reproducible way.

### Embryo injection stamp for the common house spider, *Parasteatoda tepidariorum*, opens this species to genetic manipulation

The spider injection stamp (**Fig. 2A**) generates a mold (**Fig. 2B**) that has embryo holder spaces (**Fig. 2C**). The embryos are loaded into the dish and adjusted into the spaces. They must fit deeply enough that they do not easily roll out of the spaces, but not so deep that the entire embryo fits within the space. As indicated in Materials and Methods, the mold contains 14x9 wells (126 total).

**Figure 2.**
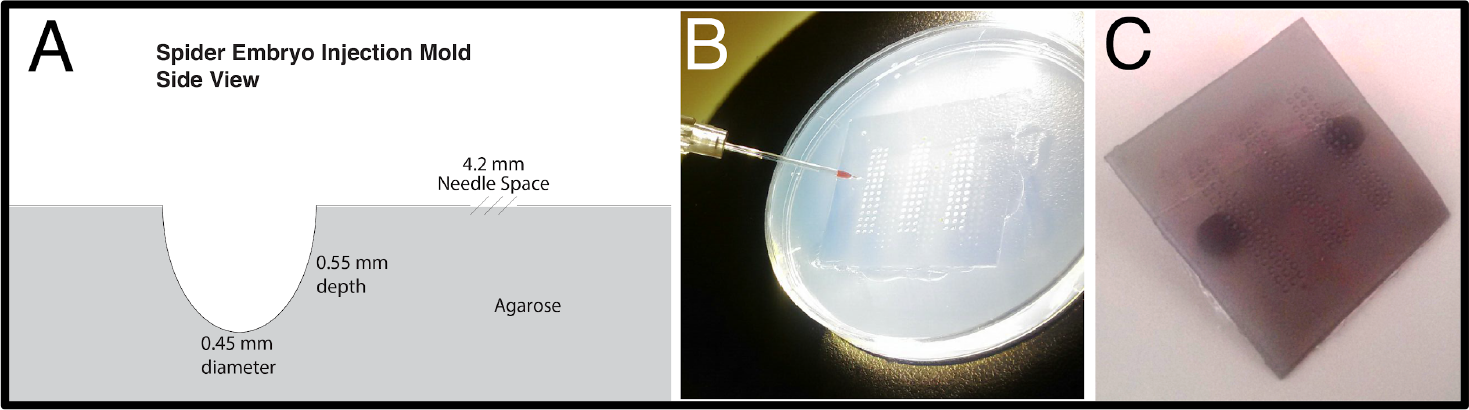
Injection stamp for *Parasteatoda tepidariorum* zygotes. **A.** Schematic of injection stamp-mold with labeled parameters. **B.** Agarose gel mold of injection stamp in a p60 petri dish with visible cup-style wells for embryos and a loaded needle prepared for injections. **C.** 3D-printed injection stamp showing the small egg-shaped protrusions that make the cups for the spider embryos.

In standard practice, spider embryonic injections were carried out on double-sided tape slides or on heptane glue coated slides. These methods allow for a survival rate at just 30% in expert hands. To hold individual embryos and avoid the sticking and toxicity of the other methods, the stamp was designed to generate wells in agarose that fit the embryos. The embryos fit snugly into the well, allowing for stability during injection, reducing the occurrence of tearing in the process. By dragging the needle against the edge of the well, the embryo can be easily slid off the needle. After injections, embryos can be easily removed from the wells with an eyelash tool, without disrupting any membrane, unlike the method involving glue, keeping the embryo intact for further analysis. In all the examined stages, both with the chorionic membrane or without, the survival rate of the embryos after injection was more than 85%.

### *Xenopus* screening stamps allows two-by-two and high-throughput comparisons even for difficult orientations

Depending on the lab, swimming tadpoles need to be anesthetized with MS-222, which after 10-15 minutes kills the animal, and screened by eye in a dish, and imaged in a hand-sculpted well in a clay dish [7]. The stamp (**Fig. 3A and 3B**) and mold (**Fig. 3C and 3D**) was designed to hold the X. *laevis* tadpole in the orientation for screening the position of its heart, laterally and dorsally/ventrally (**Fig. 3D**), and allows for side-by-side comparisons (**Fig. 3E and 3F**).

**Figure 3.**
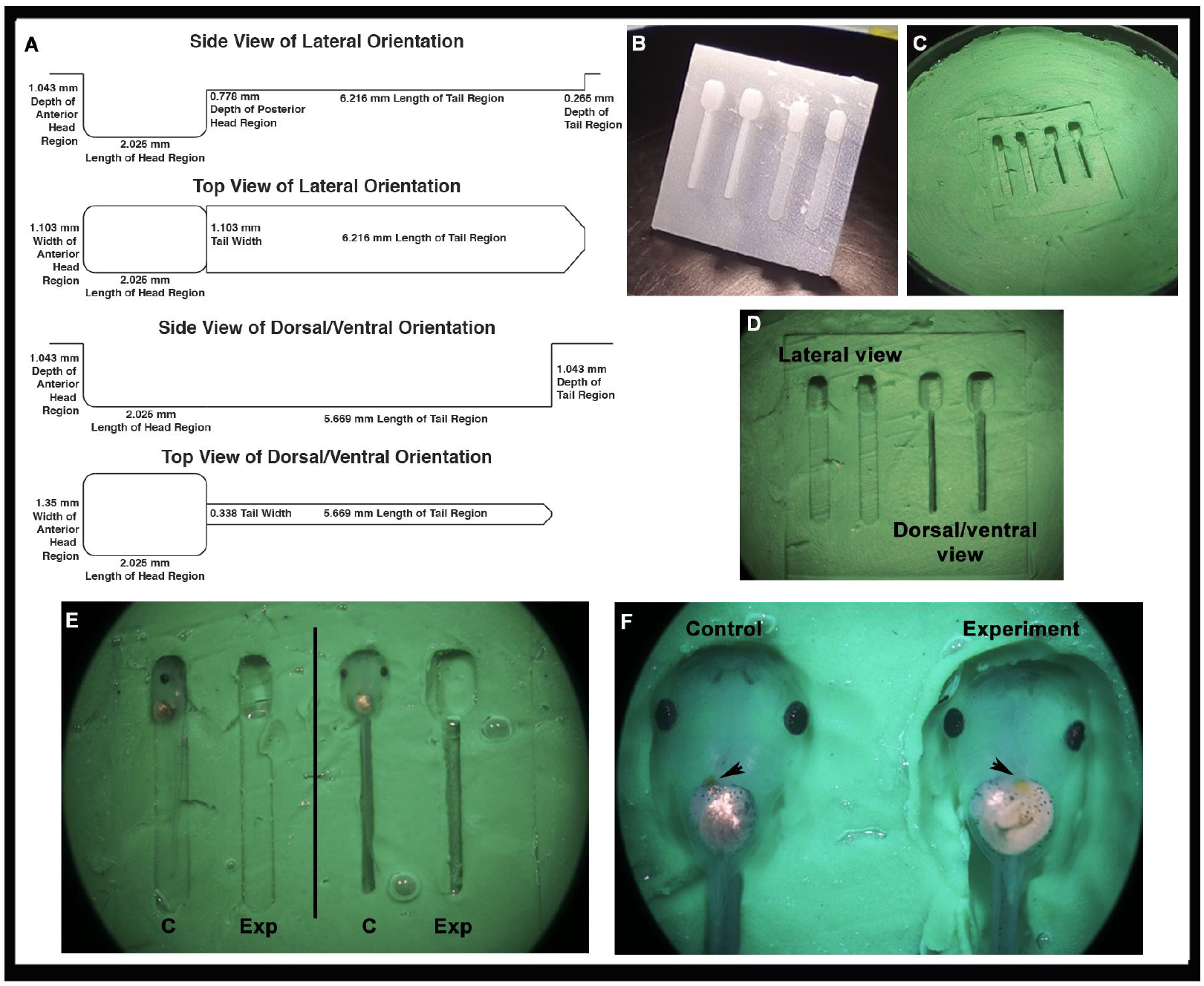
Side by side screening and imaging stamp for *Xenopus laevis* tadpoles, in stage 43. **A.** Schematic of screening stamp-mold with labeled parameters. **B.** 3D-printed screening stamp showing the lateral and dorsal/ventral extrusions. **C-D.** Clay mold for screening in a p60 petri dish. **E.** Control tadpoles orientated and arranged inside the clay mold. **F.** Detail of a two-by-two comparison between two tadpoles, ventrally orientated, which show normal and reversed heart position.

For a more high-throughput screening process, a multi-mold *X. tropicalis* tadpole stamp was designed. The stamp (**Fig. 4A and B**) and mold (**Fig. 4C**), now are designed to simultaneously screen 20 (4 x 5) tadpoles, in a drastically more efficient process. This can be optimized for any specific orientation, though in our example we designed it for screening juveniles dorsal/ventrally (**Fig. 4D**), for phenotypes expressed on only half the body, specifically in the brain (**Fig. 4E**). Tadpoles tend to lay on their side, obstructing the view of both halves of the brain, and the mold keeps them positioned properly. This method also allows for fast removal of selected individuals with a plastic eyedropper, and the next batch of tadpoles can be inserted and analyzed.

**Figure 4.**
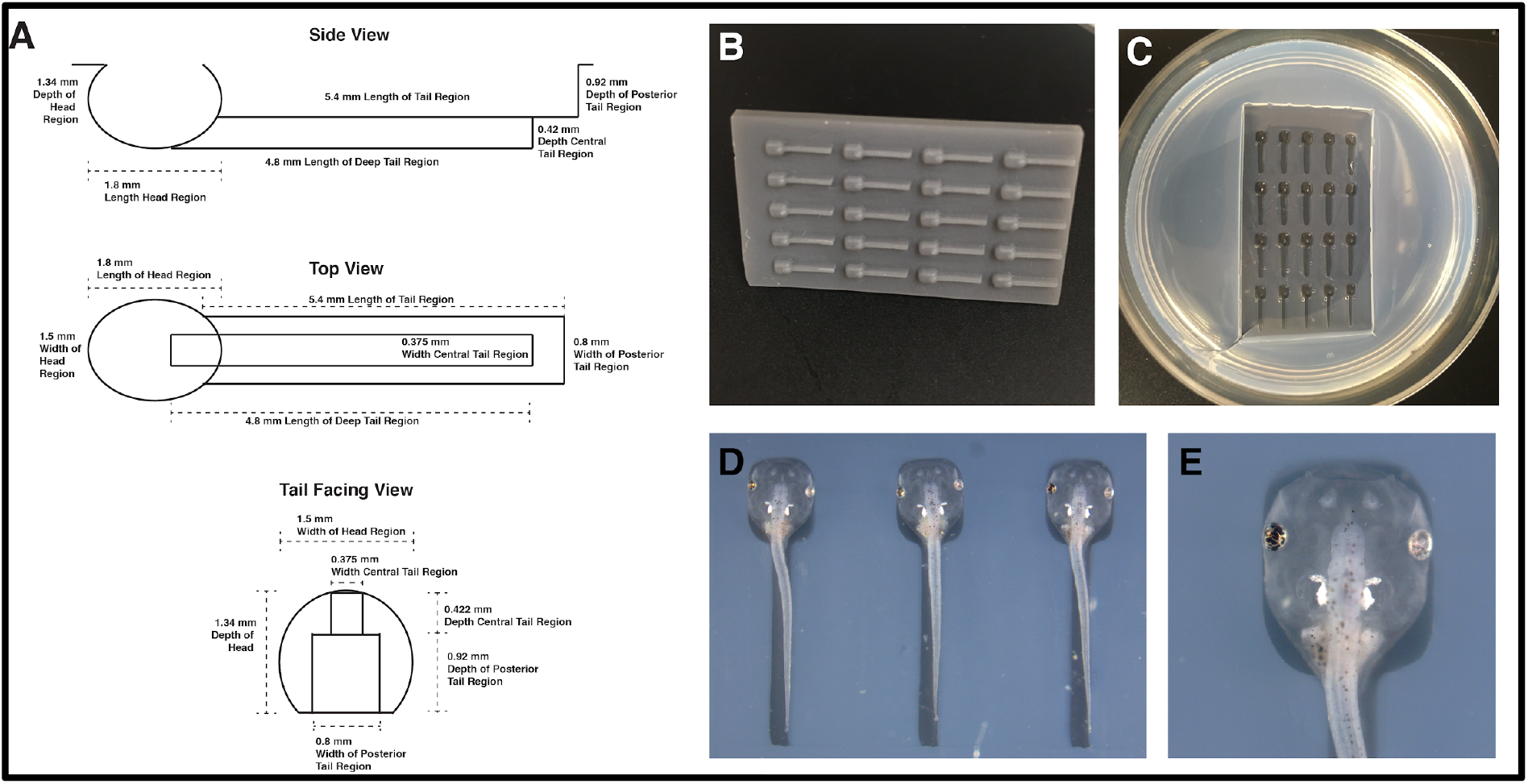
High-throughput screening and imaging stamp for *Xenopus laevis* tadpoles, in stage 43. **A.** Schematic of screening stamp-mold with labeled parameters. **B.** 3D-printed screening stamp showing the dorsal/ventral extrusions. **C.** Agarose mold for screening in a p60 petri dish. **D.** Three tadpoles orientated and arranged inside the agarose mold, ready for screening. Phenotypes on one half of the body can be easily distinguished. In this example, one eye displays abnormal development. E. Detail of a tadpole dorsally oriented with an obvious eye malformation.

### Zebrafish stamps allows several-sided, two by two sided and high-throughput confocal imaging including *in vivo*, at the same time

The stamp (**Fig. 5A and 5B**) and mold (**Fig. 5C and 5D**) was designed to hold the *Danio rerio* juveniles in a screening orientation that allows for side-by-side comparisons and confocal imaging (**Fig. 5E and 5F**). Classically, Zebrafish confocal imaging has been performed using glass-bottom dishes that allow the laser to scan the sample. A drop of low-melting temperature agarose is put on that glass surface and the zebrafish (normally up to 8 individuals) are set on the desired position with an eyelash. Then, the juveniles embedded inside the solidified agarose are imaged using a confocal. To improve this process, we maximized the size of the glass imaging surface by customizing dishes (see Materials & Methods section). This allowed the mold to orientate and stabilized up to 40 anesthetized embryos dorsal/ventrally and 40 laterally. In our example, we imaged a mutant zebrafish line for the receptor Patched, expressed along the head and the spinal cord. We can easily image all these juveniles in a lateral position, decreasing the time invested in arranging the samples. Also, the embryos can be immediately recovered since they are not embedded in agarose, and both the mold and the dish are reusable for future use.

**Figure 5.**
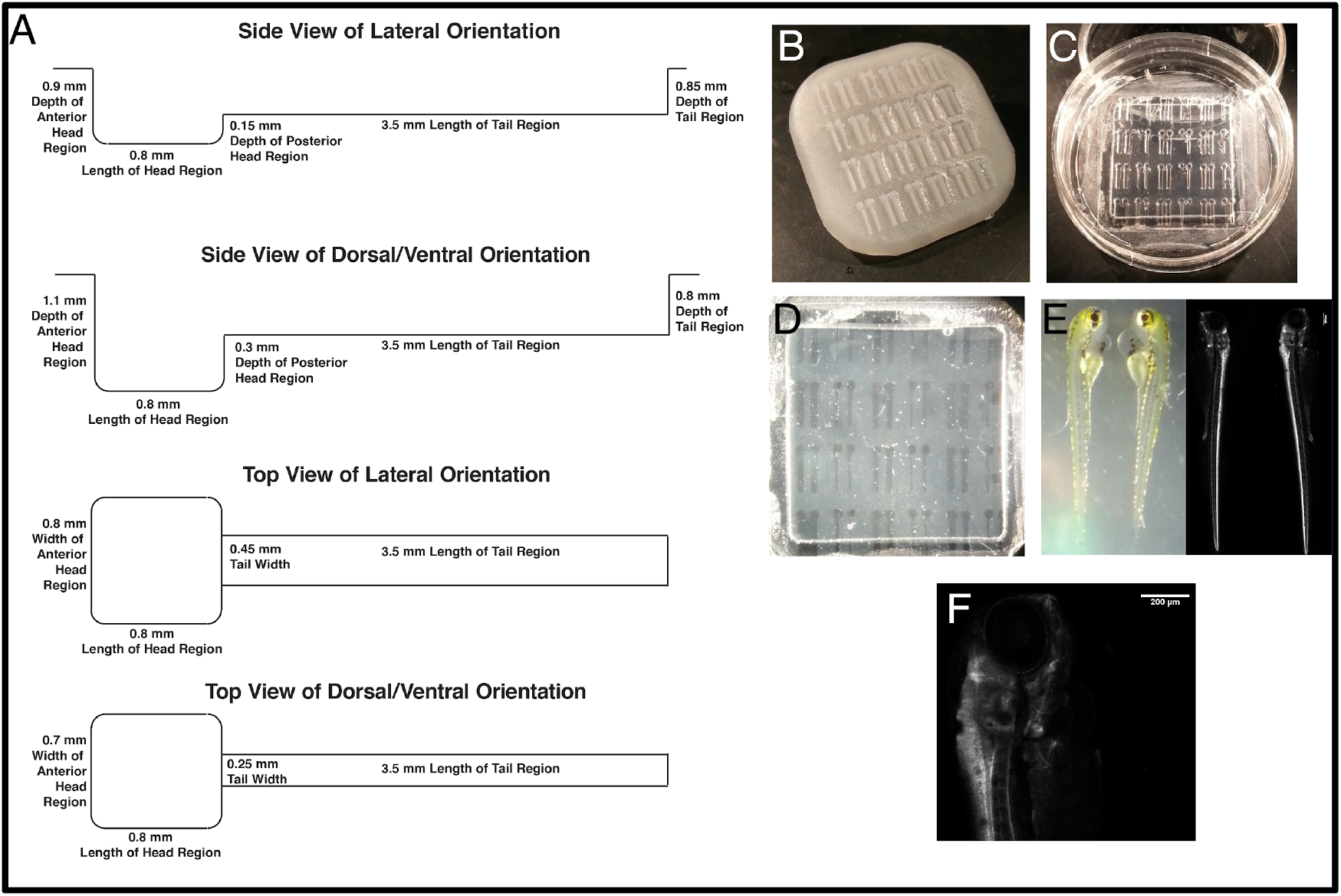
High-throughput screening and confocal imaging stamp for *Dattio rerio* juveniles, 10 dpf. **A.** Schematic of screening/imaging stamp-mold with labeled parameters. **B.** 3D-printed screening stamp showing the lateral and dorsal/ventral extrusions. **C-D.** Agarose molds for screening in a customized p35 petri dish. The bottom of the plastic dish was cut and a glass slide fixed with nail polish in order to allow the confocal laser to pass through the samples. **E.** Confocal imaging of two Zebrafish juveniles laterally oriented, which express Patched-GFP. First we imaged the juveniles in brightfield and then we used a 488 laser to image the expression of the Patched receptor, along the anterior-posterior axis. **F.** Detail of the first third of the Zebrafish juvenile located on the right side, imaged with confocal microscopy, expressing Patched-GFP.

## DISCUSSION

Ever more complex and accurate tools and techniques are required in order to keep pace with the staggeringly fast advancements in molecular biology and genetics (e.g. CRISPR/Cas9). By applying newly available mass-market 3D-printing and software technology, we have created novel stamp-mold designs for two distinct applications, though they can easily be adapted for other uses. The Form2 3D-printer is affordable when compared to other laboratory equipment, and the stamps are inexpensive and easy to replicate - once a suitable stamp is developed. Other fields already found an ally in 3D-printers; they are being used in culinary arts and the ornament and textile industries [18, 19]. In medicine, this technology already has many promising applications in prosthetics [20], for modeling tedious and dangerous surgeries before working on patients [21], and in the future we may see *in vitro* tissues printed into organs [22]. By taking advantage of this technology we were able to design helpful tools that ease embryo microinjection and screening process, especially for those without experience.

We have previously mentioned the difficulties involved in embryo injections, but more problems derive from this fact. 1) The variability due to injection number/embryo damage could affect the development, sometimes subtly, and be taken as a phenotype when in fact it is an artifact of injection, 2) the low number of embryos per experiment diminishes its statistical significance, 3) in relation to phenotypes with low penetrance (and in general), a high mortality rate hampers obtaining mutants for downstream analysis (phenotype evaluation, *in situ* hybridizations, antibody staining and western blotting, among others). Low efficiency means more injections need to be performed. Together, these issues make an already difficult task that much harder. We have shown that our tools help to ameliorate these issues.

A major hurdle to advancing the injection and screening techniques previously mentioned has been the simple fact that embryos are often rather small (100-400 μM) and commonly sticky or slippery [8, 17]. Hence, the most useful injection system would have two main traits: 1) keep the embryos confined but not stuck or compressed, so that the needle can easily penetrate the embryo without it moving, bursting, jumping or sliding; and 2) prevent the embryos from getting stuck to the needle, allowing the embryo to gently slide off the tip of the needle, avoiding any frantic movements to remove the stuck embryo, which tend to destroy it.

Embryo injections require specialized practices for any given model. For snail injections, standard practice included scratched or unscratched gelatin coated dishes in which the embryos were positioned. Separately, the standard practice for spider embryonic injections is to place the embryos on double-sided tape slides or on heptane glue coated slides. Both of these methods lead to damaged embryos, especially for novices, limiting the potential survival rate. Improving the injection process can dramatically increase the efficiency of the experiment.

We verified that these injection stamp systems not only drastically eased the injection procedure, but also decreased the mortality rate related to the manipulation of *Crepidula fornicata* and *Parasteatoda tepidariorum* embryos. Using the agarose molds drastically improved survivorship and proper development of the embryos because they are less frequently damaged and are not in toxic chemicals like heptane or the adhesive from tape. Also, the embryos are easily removed from the molds, unlike other systems that more strongly stick the embryos to a surface. The agarose molds also orient the embryos so that specific cells can be more easily injected, facilitating more specialized experiments.

After injections comes phenotype screening, often done in a juvenile stage when the organism, like larvae or tadpoles, begin to wriggle if not anesthetized or confined. Therefore, we simultaneously developed the three screening stamps to facilitate the proper orientation and quick immobilization of developing samples, allowing for fast comparisons between wild type and mutant conditions. Often phenotypes are difficult to observe, especially for beginners, and the easiest way to observe the effect of a treatment is in direct comparison to a control, which requires both organisms be placed in exactly the same orientation, in as little time as possible which the first type of screening tool achieves [Xenopus and Zebrafish imaging tool]. The second type is customized for the quick orientation and immobilization of many animals simultaneously, for high-throughput screening [Xenopus and Zebrafish high-throughput tool].

These molds facilitated the screening and imaging process, making it easier and faster to generate high quality (and aesthetically pleasing) data while critically allowing for the recovery of the *Xenopus* tadpoles from the anesthesia in all the cases. We designed a screening stamp to fit Xenopus tadpoles at stage 43, keeping in mind that they do not survive long-term exposure to anesthesia when in a fixed positioned [7]. Correctly orienting and immobilizing the embryos for phenotypic comparison becomes a countdown task, but using our screening tool makes differences between tadpoles readily apparent, crucial for fast and accurate screening under severe time pressure. It is easy to imagine how this screening stamp could also be altered to hold many more organisms for higher throughput screening. Therefore, we modified the side by side version of the *Xenopus* stamp to make it more useful for high-throughput screening of a dorsal phenotype. Similarly, we modified the *Xenopus* screening stamp to fit 10 dpf Zebrafish juveniles, while also making it compatible with confocal imaging. We showed that our system allows for simultaneous screening and imaging of five-times as many animals as the traditional methods. We were also able to perform *in vivo* recordings with our system.

A noteworthy aspect of the design is that the structure of these injection and screening molds can be easily customized to fit any embryo or larvae, simply by adjusting the parameters of the stamp, namely the width of the trough and the height of the front stop, or the diameter and depth of the embryo space, or the shape and size for the larvae - this system can be used for injecting or screening nearly any animal.

Establishing methodologies is generally an onerous process and being naïve in the technique or using a non-traditional organism for which there is no suitable method, are nearly insurmountable barriers. However, having a basic tool, a starting point, can make a big difference. Our simple stamp-mold tools and designs may facilitate the use of new organisms, while increasing the efficiency of an array of common developmental protocols.

## Acknowledgments

The authors want to thank the Marine Biological Laboratory (MBL), the Woods Hole embryology community, and the embryology course directors Richard Behringer and Alejandro Sánchez-Alvarado, for supporting the project and for providing the 3D-printer, injection systems, and the *Crepidula fornicata* adults. Thank you FormLabs, for lending the 3D-printer Form2 to the embryology course. Thank you, Detlev Arendt and the annelids module at MBL, for inspiring the snail injection tool. Thank you, Jonathan J. Henry, for sharing 3D-printing experience with us at MBL, printing some of the versions of stamps for us, and for being our expert hands injecting *Crepidula* zigotes. We thank the Henry lab, especially Kimberly Perry, for providing the *Xenopus laevis* tadpoles and their advice about their handling. Thank you, Helen Willsey, and Christian Bonato for providing us valuable feedback after using the high-throughput tadpole imaging device and the injection system for spiders, respectively, and sharing pictures for this manuscript. Thank you to Mariam Hamichi from Martín-Belmonte lab (CBMSO-Spain) and David Sánchez from Guerrero lab (CBMSO-Spain) for advising on the Zebrafish design and providing us the Zebrafish material. Finally, the authors want to thank their advisors, Cristina Grande and Lea Goentoro, for reviewing the manuscript and their advice. This research was funded by the Ministerio de Economía y Competitividad of Spain, PhD fellow BES-2012-052214 and short-term stays program EEBB-1-16-11411 to M.T.-G; and The Embryology Program at Woods Hole through the Helmsley Charitable Trust, Lorus J. & Margery J. Milne Scholarship, and The Society for Developmental Biology to M.J.A. R.M.H is a Senior Associate Dean of Biological Sciences in Berkeley and actual professor at the Embryology course at the MBL in Woods-Hole.

## Abbreviations used

3D printing: , three-dimensional printing;
dpf: , days post-fertilization;
RNAi: , ribonucleic acid interference;
CRISPR: , clustered regularly interspaced short palindromic repeats;
GFP: , green fluorescent protein;
p60 or p35: , petri dish 60 mm or 35 mm on diameter;
MS-222: , tricaine mesylate.

## References

1. Keller R & Danilchik M (1988) Regional expression, pattern and timing of convergence and extension during gastrulation of *Xenopus laevis*. Development 103: 193–209.

2. Sive HL, Grainger RM, Harland RM (2007) *Xenopus laevis* Keller Explants. Cold Spring Harb Protoc.

3. Ablain J & Zon LI (2016) Tissue-specific gene targeting using CRISPR/Cas9. Methods Cell Biol 135: 189–202.

4. Hejnol A, Martindale MQ, Henry JQ (2007) High-resolution fate map of the snail *Crepidula fornicata:* The origins of ciliary bands, nervous system, and muscular elements. Dev Biol 305(1): 63–76

5. Peyrot SM, Wallingford JB, Harland RM (2011) A revised model of *Xenopus* dorsal midline development: Differential and separable requirements for Notch and Shh signaling. Dev Biol 352(2): 254–266.

6. Price AL, Modrell MS, Hannibal RL, Patel NH (2010) Mesoderm and ectoderm lineages in the crustacean *Parhyale hawaiensis* display intra-germ layer compensation. Dev Biol 341(1): 256–266.

7. Hamilton PW & Henry JJ (2014) Prolonged *in vivo* imaging of *Xenopus laevis*. Dev Dyn 243(8): 1011–1019.

8. Henry JJ, Collins R, Perry KJ (2010) The Slipper Snail, *Crepidula:* An Emerging Lophotrochozoan Model System. Biol Bull 218(3): 211–229.

9. Kiehart DP (1982) Microinjection of echinoderm eggs: apparatus and procedures. Methods Cell Biol 25: 13–31.

10. Rombough PJ (2007) Ontogenetic changes in the toxicity and efficacy of the anaesthetic MS-222 (tricaine methanesulfonate) in zebrafish *(Danio rerio)* larvae. Comp Biochem Physiol A Mol Integr Physiol 148(2): 463–469.

11. Venero Galanternik M, Navajas Acedo J, Romero-Carvajal A, Piotrowski T (2016) Imaging collective cell migration and hair cell regeneration in the sensory lateral line. In: Detrich HW, Westerfield M, Zon LI, editors. The Zebrafish Cellular and Developmental Biology, Part B Developmental Biology. Methods in Cell Biology, Academic Press. pp. 211–256.

12. Seijo-Barandiarán I, Guerrero I, Bischoff M (2015) In Vivo Imaging of Hedgehog Transport in *Drosophila* Epithelia. In: Riobo N, editors. Hedgehog Signaling Protocols. Methods in Molecular Biology, Humana Press. pp. 9–18.

13. Hisaoka KK & Battle HI (1958) The normal developmental stages of the zebrafish, *Brachydanio rerio* (Hamilton-Buchanan). J Morphol 102: 311–327.

14. Hagedorn EJ, Ziel JW, Morrissey MA, Linden LM, Qiuyi Chi ZW, Johnson SA, Sherwood DR (2013) The netrin receptor DCC focuses invadopodia-driven basement membrane transmigration *in vivo*. J Cell Biol 201(6): 903–913.

15. Baden T, Chagas AM, Gage G, Marzullo T, Prieto-Godino LL, Euler T (2015) Open Labware: 3-D printing your own lab equipment. PLoS Biol 13(3): e1002086.

16. Wittbrodt BT, Glover AG, Laureto J, Anzalone GC, Oppliger D, Irwin JL, Pearce JM (2013) Life-cycle economic analysis of distributed manufacturing with open-source 3D printers. Mechatronics 23(6): 713–726.

17. Anderson JF (1990) The size of spider eggs and estimates of their energy content. J Arachnol 18: 73–78.

18. Gao T, Yang Z, Chen C, Li Y, Fu K, Dai J, Hitz EM, Xie H, Liu B, Song J, Yang B, Hu L (2017) Three-Dimensional Printed Thermal Regulation Textiles. ACS Nano 11(11): 11513–11520.

19. Severini C, Azzollini D, Albenzio M, Derossi A (2018) On printability, quality and nutritional properties of 3D printed cereal based snacks enriched with edible insects. Food Res Int 106: 666–676.

20. Wang S, Ghezzi CE, Gomes R, Pollard RE, Funderburgh JL, Kaplan DL (2017) *In vitro* 3D corneal tissue model with epithelium, stroma, and innervation. Biomaterials 112: 1–9.

21. Olejník P, Nosal M, Havran T, Furdova A, Cizmar M, Slabej M, Thurzo A, Vitovic P, Klvac M, Acel T, Masura J (2017) Utilisation of three-dimensional printed heart models for operative planning of complex congenital heart defects. Kardiol Pol 75(5): 495–501.

22. Pedde RD, Mirani B, Navaei A, Styan T, Wong S, Mehrali M, Thakur A, Mohtaram NK, Bayati A, Dolatshahi-Pirouz A, Nikkhah M, Willerth SM, Akbari M (2017) Emerging Biofabrication Strategies for Engineering Complex Tissue Constructs. Adv Mater 29: 1606061.

